# Transcriptomic analysis reveals that severity of infectious bursal disease in White Leghorn inbred chicken lines is associated with greater bursal inflammation *in vivo* and more rapid induction of pro-inflammatory responses in primary bursal cells stimulated *ex vivo*

**DOI:** 10.1101/2021.03.18.436042

**Authors:** Amin S. Asfor, Salik Nazki, Vishwanatha RAP Reddy, Elle Campbell, Katherine L. Dulwich, Efstathios S. Giotis, Michael A. Skinner, Andrew J. Broadbent

**Affiliations:** Birnaviruses Group, The Pirbright Institute, Ash Road, Pirbright, Woking, GU24 0NF, UK; Department of Pathology and Infectious Diseases, Faculty of Health and Medical sciences, School of Veterinary Medicine, University of Surrey, Guilford, GU2 7AL, UK; Section of Virology, Faculty of Medicine, Imperial College London, St. Mary’s Campus, Norfolk Place, London W2 1PG, UK; School of Life Sciences, University of Essex, Colchester, C04 3SQ, UK; Department of Animal and Avian Sciences, College of Agriculture and Natural Resources, University of Maryland, College Park, MD, 20742, USA

**Keywords:** IBDV, infectious bursal disease, bursa of Fabricius, inbred lines, chickens

## Abstract

In order to better understand differences in the outcome of infectious bursal disease virus (IBDV) infection, we inoculated a very virulent (vv) strain into White Leghorn chickens of inbred line W that was previously reported to experience over 24% flock mortality, and three inbred lines (15I, C.B4 and 0) that were previously reported to display no mortality. Within each experimental group, some individuals experienced more severe disease than others but line 15I birds experienced milder disease based on average clinical scores, percentage of birds with gross pathology, average bursal lesion scores and average peak bursal virus titre. RNA-Seq analysis revealed that more severe disease in line W was associated with significant up-regulation of pathways involved in inflammation, cytoskeletal regulation by Rho GTPases, nicotinic acetylcholine receptor signaling, and Wnt signaling in the bursa compared to line 15I. Primary bursal cell populations isolated from uninfected line W birds contained a significantly greater percentage of KUL01+ macrophages than cells isolated from line 15I birds (p<0.01) and, when stimulated *ex vivo* with LPS, showed more rapid up-regulation of pro-inflammatory gene expression than those from line 15I birds. We hypothesize that a more rapid induction of pro-inflammatory cytokine responses in bursal cells following IBDV infection leads to more severe disease in line W birds than in line 15I.

## Introduction

Infectious bursal disease virus (IBDV) is a member of the *Birnaviridae* family, responsible for infectious bursal disease (IBD) in chickens that is of considerable economic importance to the poultry industry worldwide. It causes morbidity and mortality in infected flocks that can be severe and is associated with welfare concerns and production losses. The virus has a preferred tropism for B lymphocytes, the majority of which reside in the bursa of Fabricius (BF). As a result, birds that recover from infection are often left immunosuppressed, less responsive to vaccination programmes against other diseases, and at an increased risk of secondary infection (1). The overall disease burden and economic impact is so extensive that IBDV has been ranked in the top 5 infectious problems of poultry (2).

For reasons that are not fully understood, the severity of clinical signs and mortality rates vary depending on the breed of the flock; for example, layer-type (LT) chickens are more susceptible to severe disease than broiler-type (BT) flocks (3, 4). There is an incentive to understand the molecular basis of IBD pathogenesis in order to breed or engineer more resistant chickens that have lower production losses due to infection. To this end, the more severe clinical signs observed in LT birds were found to be accompanied by more viral antigen in the BF, faster destruction of bursal follicles, and elevated IL-1β, IL-6 and IFN, suggesting that higher levels of pro-inflammatory and antiviral cytokines were produced in the birds more susceptible to severe disease. In addition, BT chickens had a higher number of T cells in the BF than LT birds, suggesting that the ability to mount a strong, local T-cell response during the early phase of infection correlated with reduced disease severity (3). A follow up study also found more KUL01+ macrophage-like cells accumulated in the BF of LT birds than in those of BT birds during IBDV infection, which may have been responsible for the elevated cytokine responses (4).

Genetically well-defined, inbred or partially inbred lines of chickens (5) have been maintained in the UK, at what is now the National Avian Research Facility (NARF), for over twenty generations. Mortality rates following infection with very virulent (vv) IBDV (strain CS89), were found to vary considerably between these lines, making them an attractive model to study the genetic basis for differences in disease outcome. So-called “resistant” White Leghorn lines 15I (herein referred to as 15), 7_2_, C, EB and O displayed no mortality, whereas mortality rates for White Leghorn lines N, 6_1_ and W were 6.5%, 8.3% and 24.4%, respectively, and the mortality rates for Rhode Island Red, Light Sussex and Brown Leghorn breeds were 22.2%, 40.7%, and 79.2%, respectively (6, 7). To identify gene targets that could be exploited to engineer IBD resistant chickens, several groups have conducted transcriptional profiling of the BF of the lines following IBDV infection, either by microarray (8, 9) or by RNA-sequencing (RNA-seq) (10, 11). Two studies (8, 9) compared White Leghorn line 6_1_ (8.3% mortality) with Brown Leghorn chickens (79.2% mortality) and their results indicated that an early, rapid, pro-inflammatory response and more extensive activation of apoptosis was evident in the more resistant line, suggesting that these responses may limit viral replication and pathology (8). In addition, differences in the inherent, constitutive level of expression of innate immune genes prior to infection may have affected disease outcome (9). However, the observed differences in gene expression may have been due to differences between White and Brown Leghorns, with White Leghorns mounting a more rapid inflammatory response than Brown Leghorns.

White Leghorns represent one of the leading egg producing chicken breeds of the world and there is interest in engineering more IBD-resistant birds from this breed. In order to address this, two subsequent studies examined differences in the transcriptional responses of only the inbred lines that were of the White Leghorn breed (10, 11). Birds from line P, that had more severe bursal pathology than line N, demonstrated greater up-regulation of pro-inflammatory cytokines, suggesting that excessive inflammation correlated with disease severity among the White Leghorn lines, consistent with the previous observations made in LT chickens (11). However, all the birds in these studies showed severe bursal pathology and reached their clinical humane end points by 3 days post-infection (dpi), irrespective of the line, possibly due to the birds receiving a high dose of virus (5.4 log_10_ EID_50_/bird). The authors concluded that a lower dose might provide a greater range of clinical signs and allow more differentiation between the lines (10).

Our aim was to compare the transcriptional profile of the BF between White Leghorn lines that differed in their disease outcome following infection with a lower dose (1.2 log_10_ EID_50_/bird) of vv IBDV strain UK661. We selected line W, as it had previously been reported to show the highest mortality (24.4%) (7). For comparison, we evaluated disease severity in lines 15, C, and O (12) which had shown no mortality (7). We identified genes that were differentially expressed between the lines in the BF at 3 dpi that were associated with disease severity, and we quantified the kinetics of gene expression in primary bursal cells harvested from the same lines and infected with IBDV or stimulated with LPS *ex vivo*.

## Materials and Methods

### Virus

The vv IBDV strain UK661, originally isolated in the UK from 10-day-old broilers in 1989 (13), was a kind gift from Dr Nicolas Eterradossi, ANSES, France. The virus had previously been propagated by passage in specific pathogen free (SPF) chickens followed by harvesting the BF from infected birds, homogenising the tissue in Vertrel XF (Sigma-Aldrich) and harvesting the aqueous phase. Virus was titrated in embryonated chicken eggs by inoculation on to the chorioallantoic membrane. Embryos (7 dpi) were inspected for clinical signs associated with IBDV infection and the titre of the virus was quantified by egg infectious dose -50 (EID_50_) (1). The virus was also titrated in the immortalised B cell-line DT40 (14). At 3 dpi, wells were fixed in 4% paraformaldehyde and stained with a mouse monoclonal antibody against the IBDV VP3 protein (15) that co-localises with IBDV virus factories (16) and a goat anti-mouse secondary antibody conjugated to Alexa-Fluor 488 (Invitrogen). Wells were scored as positive or negative based on the presence of infected cells, and the titre was expressed as tissue-culture infectious dose-50 (TCID_50_) (17).

### Chickens

SPF chickens of inbred lines W, 15, C, and O were obtained from unvaccinated flocks, supplied by the National Avian Research Facility (NARF), UK. Inbred lines were all of the White Leghorn breed and had been maintained for over 20 generations by full sibling mating. The inbred lines had the following MHC haplotypes: B^14^ (W), B^15-^(15), B^4^/B^4^ (C), and B^21^ (O). More details of the lines can be found at http://www.narf.ac.uk/chickens/. Infected birds and age-matched mock-inoculated control birds were housed in separate experimental animal rooms to prevent contamination.

### Inoculation of chickens with IBDV

Six birds from each inbred line (n=24) were infected at 2-3 weeks of age with 1.2 log_10_ EID_50_ of UK661 vvIBDV strain via the intra-nasal route. Virus was diluted in sterile phosphate buffered saline (PBS) and 100μL was administered to each bird, 50μL per nares. Additionally, six birds from each inbred line (n=24) were mock infected with PBS alone. Birds were weighed daily and monitored for clinical signs at least twice daily. Clinical scores were recorded according to a points-based scoring system developed at The Pirbright Institute (18) that characterized disease as mild (1–7), moderate (8–11), or severe (12–17). Briefly, birds were scored on their appearance, behavior with and without provocation and handling, and included an assessment of the combs, wattles, feathers, posture, eyes, breathing, interactions with the rest of the flock, ability to evade capture, and crop palpation (18). Birds were humanely culled when they reached humane endpoints of 12 or above. Three days post-inoculation, the birds were humanely culled by cervical dislocation and subject to post-mortem (PM) analysis. Pectoral and thigh muscles were inspected for the presence of petechial haemorrhages, spleens were inspected for enlargement and discolouration, and bursal tissue was inspected for evidence of congestion, oedema, or haemorrhage. The BF was divided into two sections, one stored in RNALater (Thermo Fisher Scientific) for RNA extraction and quantification of gene expression by RTqPCR and RNA-Seq, and one frozen in tissue freezing medium (Leica, Australia) for cryosectioning. All animal procedures were conducted following the approval of the Animal Welfare and Ethical Review Board (AWERB) at The Pirbright Institute, under Home Office Establishment, Personal and Project licenses, and conformed to the United Kingdom Animal (Scientific Procedures) Act (ASPA) 1986.

### Pathological scoring of bursal tissue

Bursal samples were sectioned using a CM1860 UV Cryostat Ag Protect machine (Leica), mounted onto glass slides and stained with a mouse monoclonal antibody against the IBDV VP3 protein and a goat-anti-mouse secondary antibody conjugated to Alexa-Fluor 488. Tissue sections were stained with DAPI before ProLong Glass antifade (Invitrogen) and a coverslip were added. Bursal tissue was observed microscopically using a TCS SP5 confocal microscope (Leica), and evaluated based on the severity of follicle damage, necrosis and lymphocyte depletion using a scoring system that was previously described by (19).

### Isolation of primary bursal cells

Primary bursal cells were obtained from the BF of line 15 and line W birds as previously described (20, 21). Briefly, the BF was harvested aseptically post-mortem and washed in sterile PBS. An enzyme solution containing 2.2 mg/mL of collagenase D (Sigma-Aldrich) was used to digest the bursal tissue. Once digested, the tissue was passed through a 100μm Falcon cell strainer (Thermo Fisher Scientific) into cell medium containing HBSS (Gibco) supplemented with 7.5% sodium carbonate and 500mM EDTA (Sigma-Aldrich), before pelleting. Cells were re-suspended in RPMI before undergoing centrifugation at 2,000 rpm for 20 min at 4°C over Histopaque 1077 (Sigma-Aldrich). Bursal cells that banded at the interface of the medium and histopaque were collected, washed in PBS, counted, and re-suspended in Iscove’ s modified Dulbecco’ s medium (IMDM) supplemented with FBS, chicken serum (Sigma-Aldrich), insulin transferrin selenium (Gibco), beta-mercaptoethanol (Gibco), penicillin, streptomycin, nystatin and chCD40L (20, 21) (complete B cell media).

### Infection of primary bursal cells with IBDV

Primary bursal cells were pelleted and re-suspended in 1mL of IBDV diluted in complete B cell media at an MOI 3 for 1 hour at 37°C, after which the cells were washed and re-suspended in complete B-cell media at a concentration of 1×10 ^7^ cells per mL and cultured in 24-well plates at 37°C, 5% CO_2_ until 3, 6, and 18 hours post infection (hpi), whereupon RNA was extracted as described.

### Stimulation of primary bursal cells with Lipopolysaccharide (LPS)

Primary bursal cells were pelleted and re-suspended in 1mL of LPS (Sigma-Adrich) diluted to a concentration of 100ng/mL in complete B-cell media for 1 hour at 37°C, after which the cells were washed and re-suspended in complete B-cell media at a concentration of 1×10 ^7^ cells per mL and cultured in 24-well plates at 37°C, 5% CO_2_ until 3, 6, and 18 hpi, whereupon RNA was extracted as described.

### RNA extraction

Briefly, either primary bursal cells were re-suspended in RLT buffer, or 20-30mg of bursal tissue was homogenised in RLT buffer, and RNA was extracted from using an RNeasy kit (Qiagen) according to the manufacturer’ s instructions. RNA in the samples was quantified using a Nanodrop Spectrophotometer (Thermo Scientific) and checked for quality using a 2100 Bioanalyzer (Agilent Technologies). All RNA samples used in RNA-seq had an RNA integrity number (RIN) from 7 to 9.4.

### Quantification of gene expression by RTqPCR

RNA samples were reverse transcribed using SuperScript III Reverse Transcriptase (Thermo Fisher Scientific) and a random primer, according to the manufacturer’ s instructions, and gene expression was quantified by qPCR using primers that are listed in table S5. Briefly, the template and primers were added to a Luna Universal qPCR Master Mix (New England BioLabs) and reactions were performed on a 7500 Fast Real-Time PCR system (Life Technologies) according to the manufacturer’ s instructions. Virus and host-cell gene expression was normalized to the housekeeping gene RPLPO and expressed relative to mock controls in a comparative ΔΔCT method.

### Preparation of cDNA libraries and RNA sequencing

cDNA libraries were constructed by the Beijing Genomics Institute (BGI). Briefly, poly-A containing mRNA molecules were purified using poly-T oligo-attached magnetic beads (NEB) that were subsequently fragmented into shorter mRNA fragments using divalent cations under an elevated temperature (RNA fragmentation reagents kit, Ambion). Reverse transcriptase and a random primer (Invitrogen) were then used to synthesize first strand cDNA from the cleaved RNA fragments. Second strand cDNA synthesis proceeded with a DNA Polymerase I (NEB) and RNase H (Invitrogen), and the RNA template was removed. An ‘ A’ base was added to the resulting cDNA fragments, that was subsequently ligated to an adapter. Finally, the products were enriched by PCR before purification with the MinElute PCR Purification Kit (Qiagen). This purified cDNA library was used in RNA-Seq analysis. RNA-Seq reads were generated was by BGI with the Illumina HiSeq2500 platform. Following sequencing, initial analysis was conducted by BGI using SOAPnuke software (v1.5.5) to filter the sequence reads with the following parameters: -n 0.1 -l 20 -q 0.4 -i -A 0.25 -Q 2 -G --seqType 1.Reads with adaptors were removed, as were reads with unknown bases at a frequency of >0.1%. Reads comprising over 40% of bases with a quality score for each individual base of less than 20, were determined as low quality and these reads were removed. The remaining reads (over 70 million per sample) were defined as ‘ clean reads’ and stored in FASTQ format. FASTQ files were imported into CLC Bio’ s Genomics Workbench (CLC Bio, Qiagen Bioinformatics, Aarhus, Denmark) v12.0.2 for quality-control processing and analysis. RNA-Seq data are deposited in the GEO and SRA archives at NCBI (Accession number GSE166026).

### RNA-seq read mapping and differentially expressed gene (DEG) analysis

RNA-seq read mapping and DEG analysis was performed as previously described (22, 23). Briefly, RNA-seq reads were subjected to quality trimming before mapping to the ENSEMBL galGal5-annotated assembly (GRCg6a; 03-07-2019) for quantitative analysis of expression. The fold change and False Discovery Rates (Bonferroni) were calculated using CLC Bio’ s RNA-Seq Analysis tool (v2.18), and differential expression within the RNA-Seq data was analysed using CLC Bio’ s Differential Expression for RNA-Seq tool (v2.2). Two comparisons were made in this study: line 15 versus line W and high versus low clinical score. For line 15 versus W, the gene expression in the BF samples from line 15 birds infected with UK661 at 3 dpi were compared to the gene expression in the BF samples from line W birds infected with UK661 at 3 dpi, using line W as a baseline. For high versus low clinical score, the gene expression was determined in the BF samples from birds that had a clinical score of 8-14 and compared to birds that had a clinical score of 1-5. For both high and low clinical scores, we selected birds for which there was no significant difference in viral replication. Birds within the high score group were comprised of one line W, one line 15, two line C, and two line O (n = 6), and birds within the low score group were comprised of one line W, one line 15, two line C, and two line O (n = 6). Data mining, gene ontology (GO) enrichment analysis, and pathway analysis was conducted using PANTHER (Protein Analysis Through Evolutionary Relationships) Version 14.1 (released 2019-03-12).

### Flow cytometry

Primary cells were isolated from the BF of an additional five uninfected line W and an additional five uninfected line 15 birds as described. Cells were re-suspended in FACS buffer (1% bovine serum albumin (BSA) in 1x PBS), plated in U-bottom 96-well plates and treated with 4% BSA for 20 min at room temperature (RT) to block the Fc receptors. Cells were then surface stained with mouse anti-chicken Bu-1-FITC (Clone AV20; Southern Biotech, Birmingham, AL, USA) and mouse anti-chicken monocyte/macrophage-PE (Clone KUL01, Southern Biotech, Birmingham, AL, USA) antibodies in the dark for 20 min at RT. Finally, the cells were fixed in 4% paraformaldehyde, re-suspended in 400 µL FACS buffer and subject to flow cytometry using BD LSRFortessa™ Flow cytometer (BD Biosciences). Ten thousand events were collected per sample, and gated based on forward scatter (FSC) and side scatter (SSC). Data were analysed using FlowJo software (FlowJo_v_10.6.2) to count the percentage of Bu1+ and KUL01+ cells in the BF population.

### Statistical analyses

All statistics in this study, unless otherwise stated, were performed using the statistical function on GraphPad Prism v7 and consisted of a one-way ANOVA with a Tukey’ s multiple comparison test, following a Shapiro-Wilk normality test to check the normal distribution of data sets. For gene enrichment analyses, significance was determined using PANTHER. Values <0.05 were considered significant (*p<0.05, **p<0.01,***p<0.001, ****p<0.0001).

## Results

### Clinical signs in experimentally inoculated inbred lines

Mock-infected birds increased weight from 100% on 0 dpi to an average of 118% (W), 120% (15), 111% (C), and 118% (O) on 3 dpi (Figure 1A). In contrast, birds infected with IBDV strain UK661 gained significantly less weight, increasing to an average of only 112% (W) (p<0.05), 111% (15) (p<0.05), 104% (C) (p<0.01), and 105% (O) (p<0.01) by 3 dpi. All infected birds showed signs of disease at 3 dpi, with average clinical scores of 4.67 (W), 3.17 (15), 5.67 (C) and 4.83 (O) (Figure 1B), although there was a wide range of disease severity in every infected group. The number of birds reaching a clinical score of 11 or above (the humane end points of the study) was 1/6 (17%) from line W, 1/6 (17%) from line 15, 2/6 (33%) from line C and 2/6 (33%) from line O (Figure 1B). Taken together, these data demonstrate that birds from all inbred lines were susceptible to IBD, but more line 15 birds experienced milder disease than the other lines.

**Figure 1.**
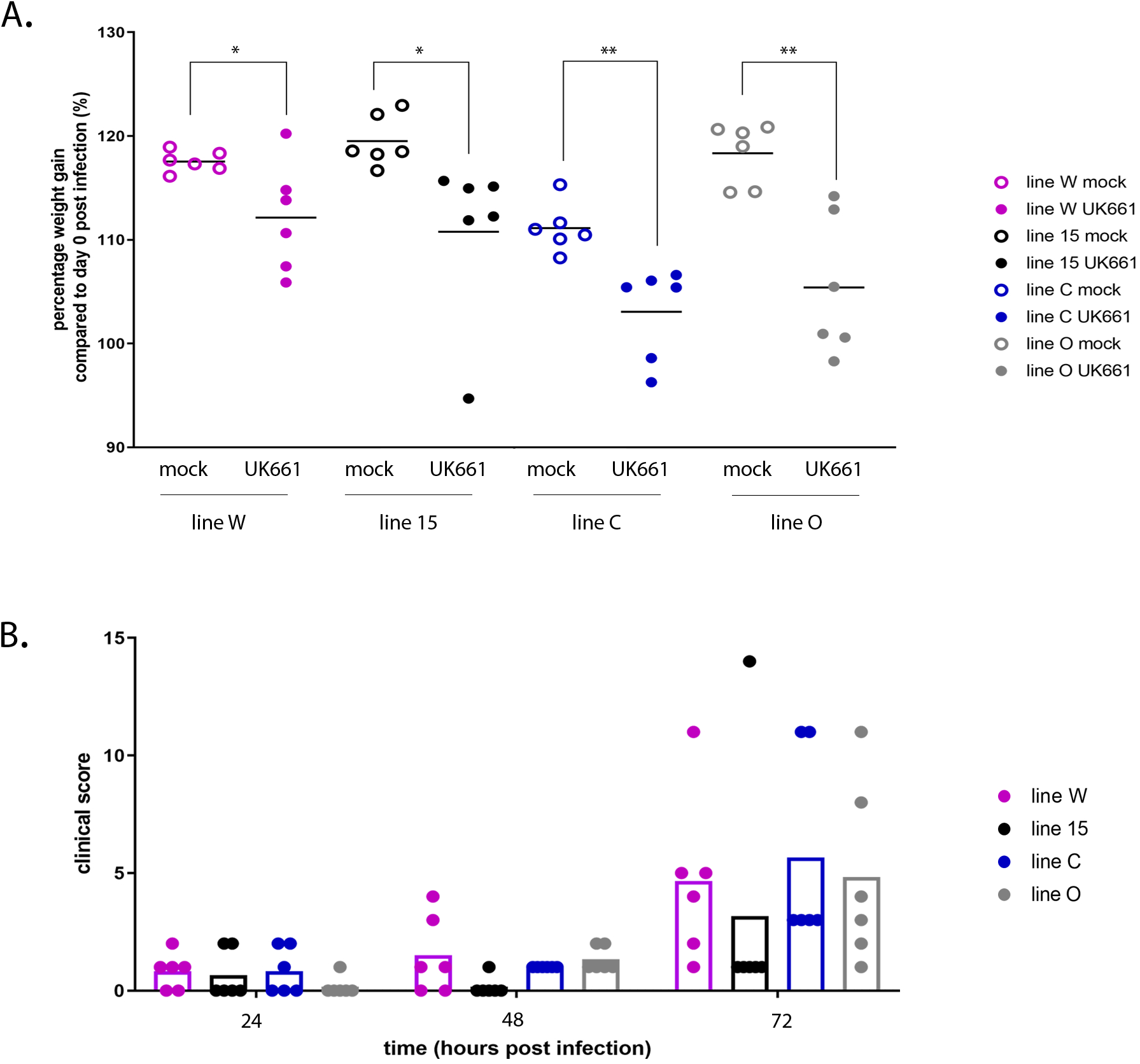
IBDV strain UK661 caused disease in all experimentally inoculated inbred lines with varying severity, but line 15 had milder clinical signs. Six birds at 2-3 weeks of age from each of lines W, 15, C and O were inoculated with 1.2 log_10_ EID_50_ of IBDV strain UK661 and six birds from each of these lines were mock-inoculated with PBS intranasally. Birds were weighed daily and the percentage weight gained at 3 dpi relative to the day of inoculation (day 0) was plotted for mock-infected birds (open circles) and IBDV-infected birds (closed circles) from lines W, 15, C and O (A). The clinical score was determined daily based on an in-house scoring system and plotted for each bird. Each dot represents one bird and each bar represents the average clinical score for the indicated line (B).

### Gross-pathology, histo-pathology and viral replication in experimentally inoculated inbred lines

Carcasses were inspected for evidence of gross pathology, including oedema, congestion and haemorrhage in the BF, enlargement and discolouration of the spleen and haemorrhage in the pectoral and thigh muscles. Mock-inoculated control chickens had no gross abnormalities on post-mortem. In contrast, the percentage of infected carcasses that had evidence of gross pathology in the BF, spleen, or muscle was 3/6 (50%) in line W, 2/6 (33%) in line 15, 6/6 (100%) in line C and 4/6 (67%) in line O (Figures 2A and 2B), but there was no statistically significant difference in the BF weight: Body weight (BF:BW) ratio between mock and infected groups for any of the lines (Figure 2C). The bursa from each bird was cryosectioned, imaged by immunofluorescence confocal microscopy (Figure 3 A-E), and the lesion score was determined. Every mock-inoculated bird had a post-mortem bursal lesion score of 0. In contrast, the average bursal lesion scores for infected birds were 2.6 (W), 1.3 (15), 2.4 (C) and 3.2 (O) (Figure 3F). The bursal lesion score correlated with the clinical scores, for example, in line 15 birds, the individual with the highest bursal score (5) also had the highest clinical score (14). The average fold change in viral RNA copy number was significantly lower in line 15 birds (6.3 log_10_) compared to lines C or O (7.0 and 6.8 log_10_, respectively) (p<0.01) (Figure 3G). Taken together, these data suggest that birds from line 15 had less gross and histo-pathology than the other lines in the study and less IBDV replicating in the BF than lines C or O.

**Figure 2.**
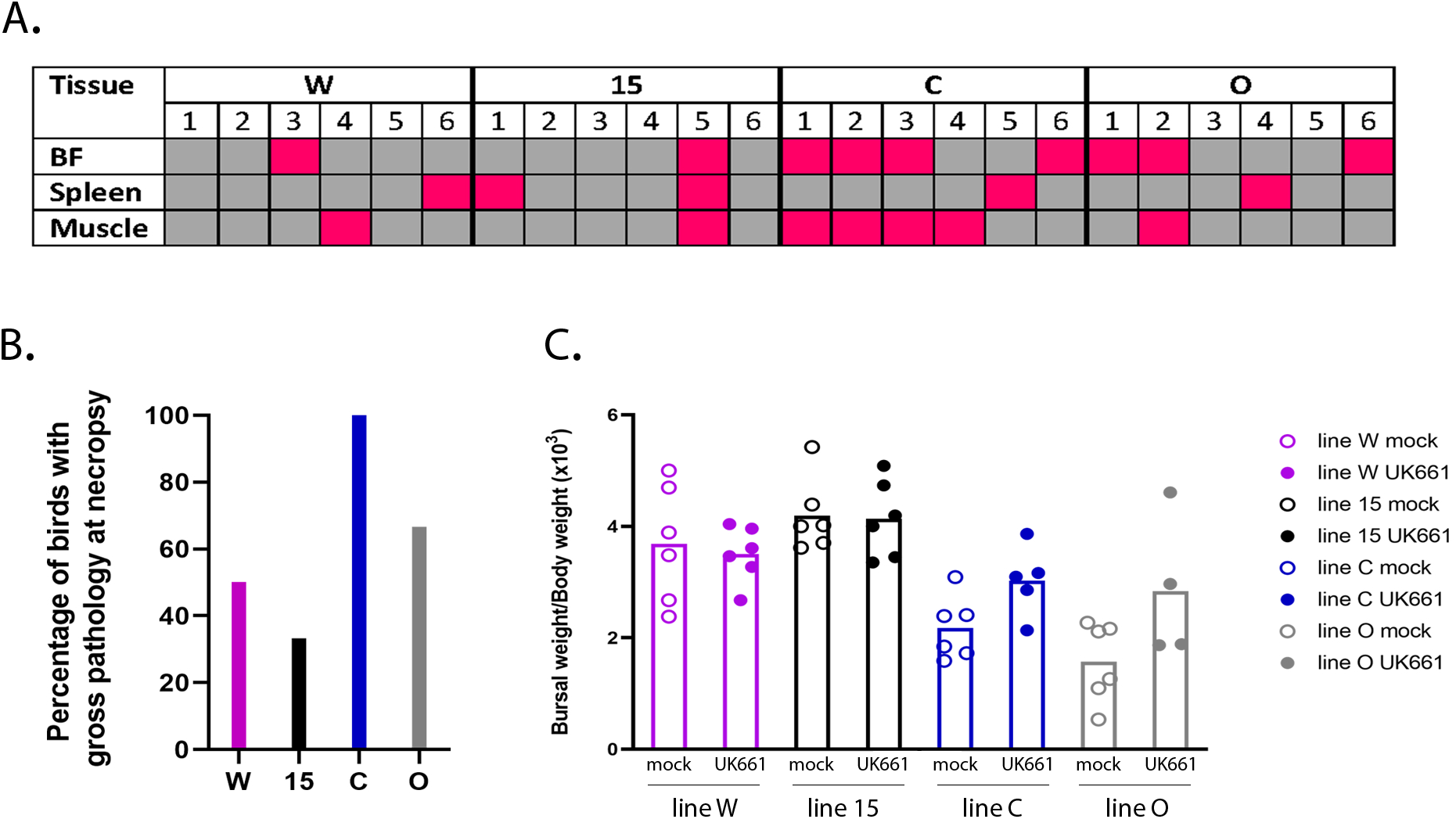
Line 15 had less gross pathology than the other lines in the study. Carcasses were inspected for evidence of gross pathology, including oedema, congestion and haemorrhage in the BF, enlargement and discolouration of the spleen, and haemorrhage in the pectoral and thigh muscles in 6 infected birds from each of line W, 15, C and O at 3 days post infection. Pink represents detection of pathology; grey represents lack of detected pathology (A). The percentage of infected birds from line W, 15, C and O showing gross detected pathology was plotted (B). The BF weight was determined at necropsy and expressed relative to body weight BF weight: body weight ratio *100) for lines W, 15, C and O in mock and UK661 infected birds and plotted. Each dot represents one bird and each bar represents the average for the indicated line (C).

**Figure 3.**
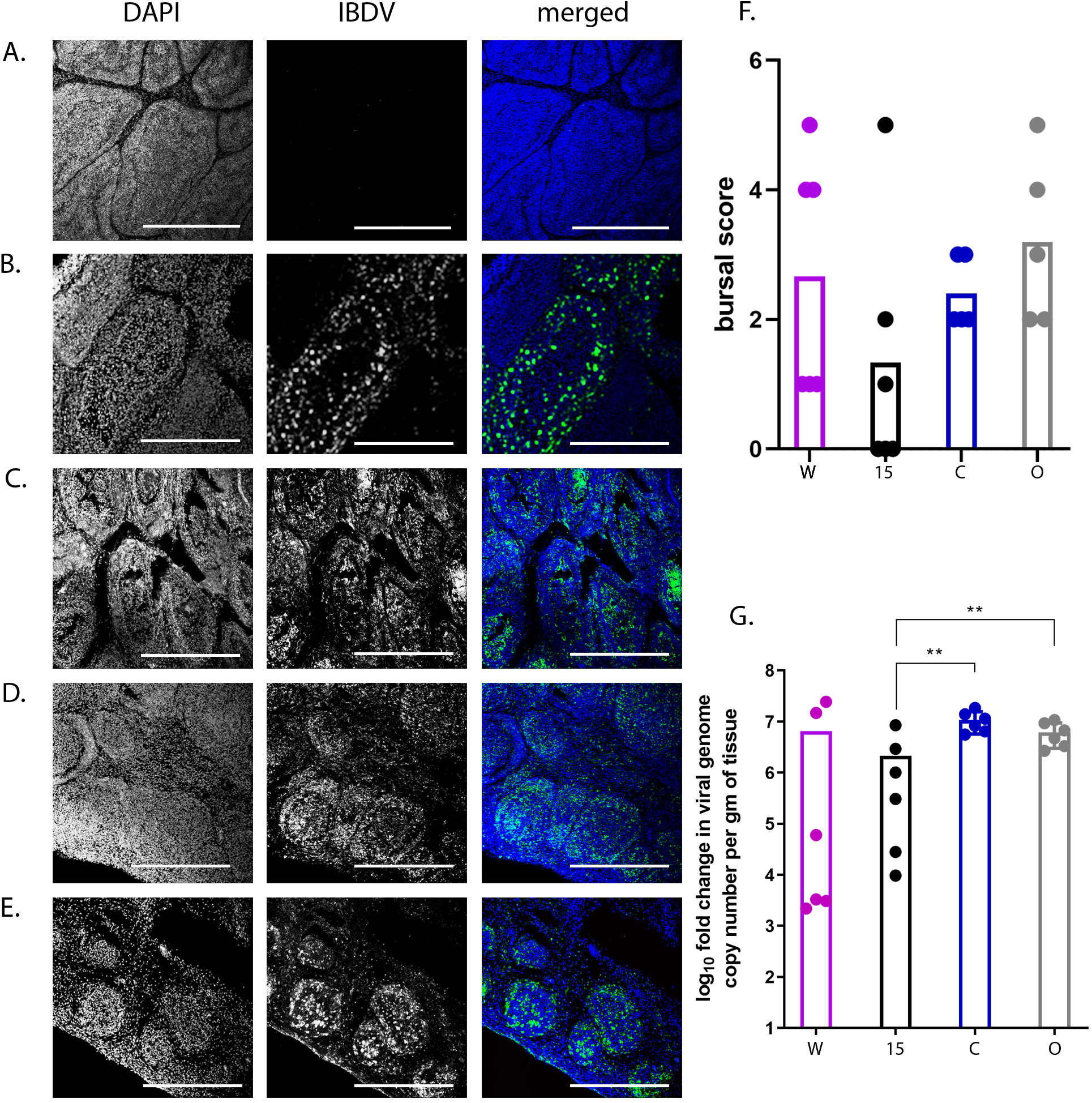
Line 15 had less histopathology and lower levels of viral replication than other lines in the study. Bursal samples were harvested from birds at 3 dpi, frozen in tissue freezing medium and cryo-sectioned. Tissue sections were stained with a mouse anti-IBDV VP3 monoclonal IgG antibody and a goat-anti-mouse IgG secondary antibody conjugated to AlexaFluor 488 (green) and cell nuclei were stained with DAPI (blue) and imaged by confocal microscopy. Scale bars represent 250μm (A-E). Sections were imaged from mock-inoculated and UK661 infected line W birds (A and B), and UK661 infected line 15 (C), line C (D) and line O (E) birds. The extent of damage to each BF was scored according to published protocols (19). The lesion scores of the images were 0, 4, 5, 2 and 5, respectively. BF lesion scores were plotted for birds of line W, 15, C and O. Each dot represents one bird and each bar represents the average for the indicated line (F). RNA was extracted from the BF samples, reverse transcribed to cDNA, and amplified by qPCR using primers specific to a virus gene (VP2) and a housekeeping gene (RPLPO). The fold change in viral RNA was determined for each bird, normalised to RPLPO, and expressed per g of BF tissue relative to mock infected control samples in a ΔΔCT analysis.The Log_10_ fold change of viral VP2 copy number per g of BF tissue was plotted for birds of line W, 15, C and O (G). Each dot represents one bird and each bar represents the average for the indicated line. Data passed Shapiro-Wilk normality tests before being analysed by one-way ANOVA and Tukey’ s multiple comparison tests (**p<0.01).

### Transcriptomic analysis of the BF reveals candidate genes associated with more severe disease

The transcriptional profile of the BF was compared between six infected birds with a clinical score of 8-14 (“high clinical score”) and six infected birds with a clinical score of 1-5 (“low clinical score”) at 3 dpi, with low clinical score as a baseline (Figure 4A). There was no significant difference in the degree of viral replication between the two groups (p = 0.08) (Figure 4B). RNA was extracted from the BF of the birds and subject to RNA-seq. A total of 475 Differentially expressed genes (DEGs) were identified (Tables S1 and S2), of which 86 were found to be up-regulated in the high clinical score group compared to the low clinical score group (Table S1 and Figure 4C). Up-regulated genes associated with pathways involved in inflammation mediated by chemokine and cytokine signaling, cytoskeletal regulation by Rho GTPases, nicotinic acetylcholine receptor signaling, and Wnt signaling were over-represented compared to other pathways (Figure 4D). Genes that were up-regulated in these pathways included: MYLK2 (47 fold), MYH1A (967 fold), MYH1B (475 fold, MYH1C (151 fold), MYH1D (42 fold), MYH1F (240 fold), MYH7 (5 fold), MY7B (12 fold), PTGS1 (5 fold), CHRND (147 fold) and CHRNG (86 fold). The transcriptional profile in the BF was also compared between infected line W and line 15 birds. A total of 151 DEGs were identified (Tables S3 and S4), of which 120 were significantly down-regulated in line 15 birds compared to line W (i.e. were up-regulated in line W compared to line 15) (Table S4 and Figure 5A). Interestingly, the same pathways were also over-represented, namely those associated with inflammation mediated by chemokine and cytokine signaling, cytoskeletal regulation by Rho GTPases, nicotinic acetylcholine receptor signaling, and Wnt signaling (Figure 5B). Genes that were down-regulated in line 15 compared to line W included: PLCH1 (6 fold), ACTA1 (15 fold), ACTC1 (25 fold), MYLK2 (87 fold), MYH1A (1589 fold), MYH1B (716 fold), MYH1C (270 fold), MYH1 F (263 fold), MYO3B (5 fold), and CHRND (175 fold). A total of 53 (62%) of genes that were up-regulated in birds with a high clinical score compared to low clinical score were also down-regulated in line 15 birds (up-regulated in line W birds) (Figure 5C), and a total of 4 (1%) of genes that were down-regulated in birds with a high clinical score compared to low clinical score were also up-regulated in line 15 birds (down-regulated in line W birds) (Figure 5D). Taken together, these data indicate that birds that experienced more severe disease induced more inflammation and tissue remodelling related transcripts in the BF than birds that experienced mild disease.

**Figure 4.**
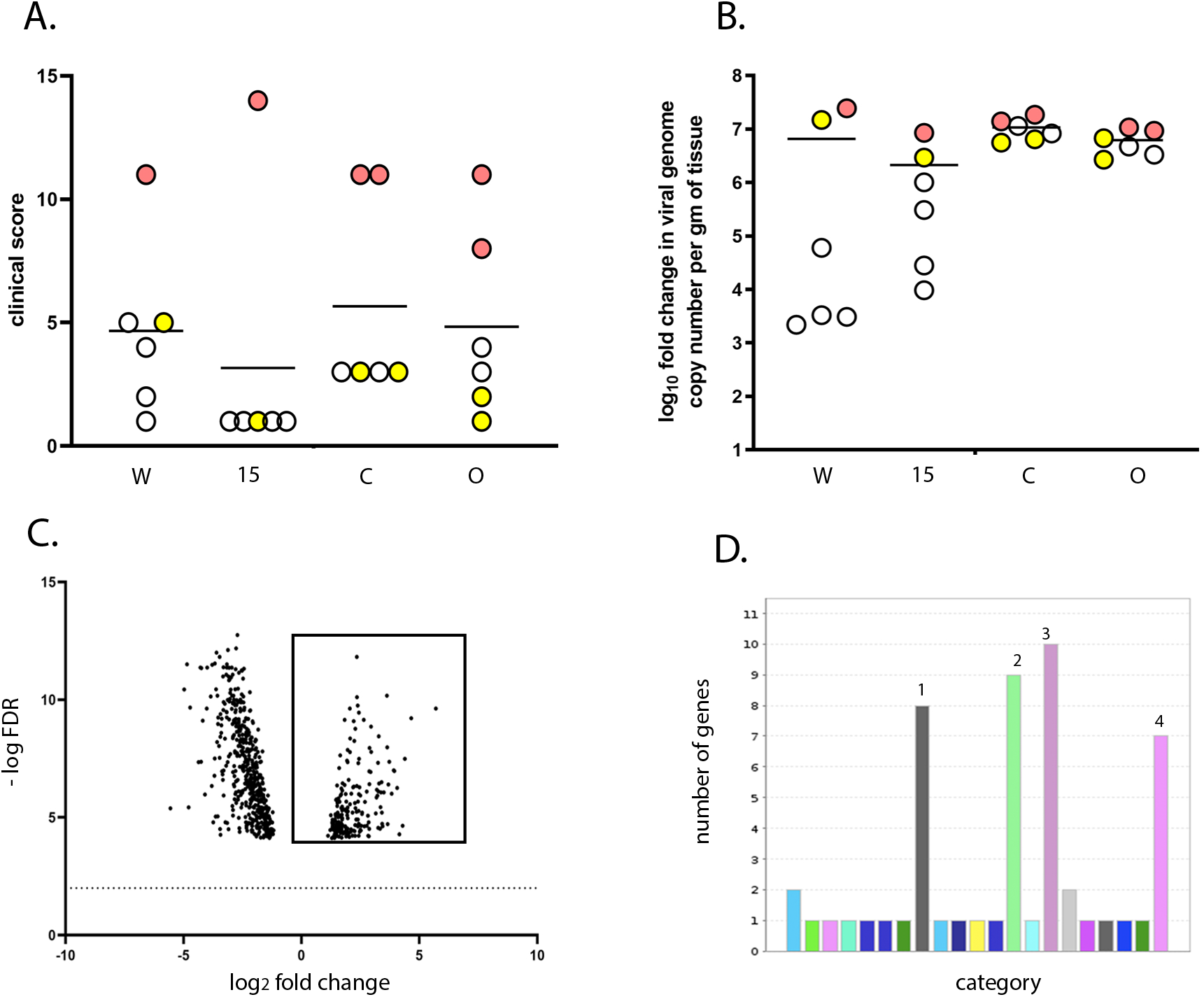
Birds with a high clinical score showed up-regulation of pathways involved in inflammation mediated by chemokine and cytokine signaling, cytoskeletal regulation by Rho GTPases, nicotinic acetylcholine receptor signaling, and Wnt signaling in the BF compared to birds with a low clinical score. The transcriptional profile of 6 birds with a “high clinical score” (8-14; pink dots) was compared with that of 6 birds with a “low clinical score” (1-5; yellow dots) at 3 dpi, with low clinical score as a baseline. Both the high and low clinical score groups consisted of one bird from line W, one from line 15, two from line C and two from line O (A), and there was no significant difference in the average viral replication between the two groups (B). RNA was extracted from the BF of the birds and subjected to RNA-seq. The expression of the target genes was reported as fold-change (FC), with values ≥ 2 considered as up-regulated, and values ≤ -2 considered as down-regulated. The false discovery rate (FDR) was calculated and a volcano plot of –log FDR over log_2_ FC plotted; FDR values of ≤0.05 were considered as significant. Each dot represents one gene and genes meeting the cut-off criteria for FC and FDR, therefore considered as up-regulated in birds with a high clinical score compared to a low clinical score are boxed (C). Pathway analysis of the selected genes with Protein ANalysis THrough Evolutionary Relationships (PANTHER) revealed 4 pathways that were significantly up regulated: 1 (grey)-inflammation mediated by chemokine and cytokine signaling, 2 (green)-cytoskeletal regulation by Rho GTPases, 3 (purple)-nicotinic acetylcholine receptor signaling, and 4 (pink)-Wnt signaling (D).

**Figure 5.**
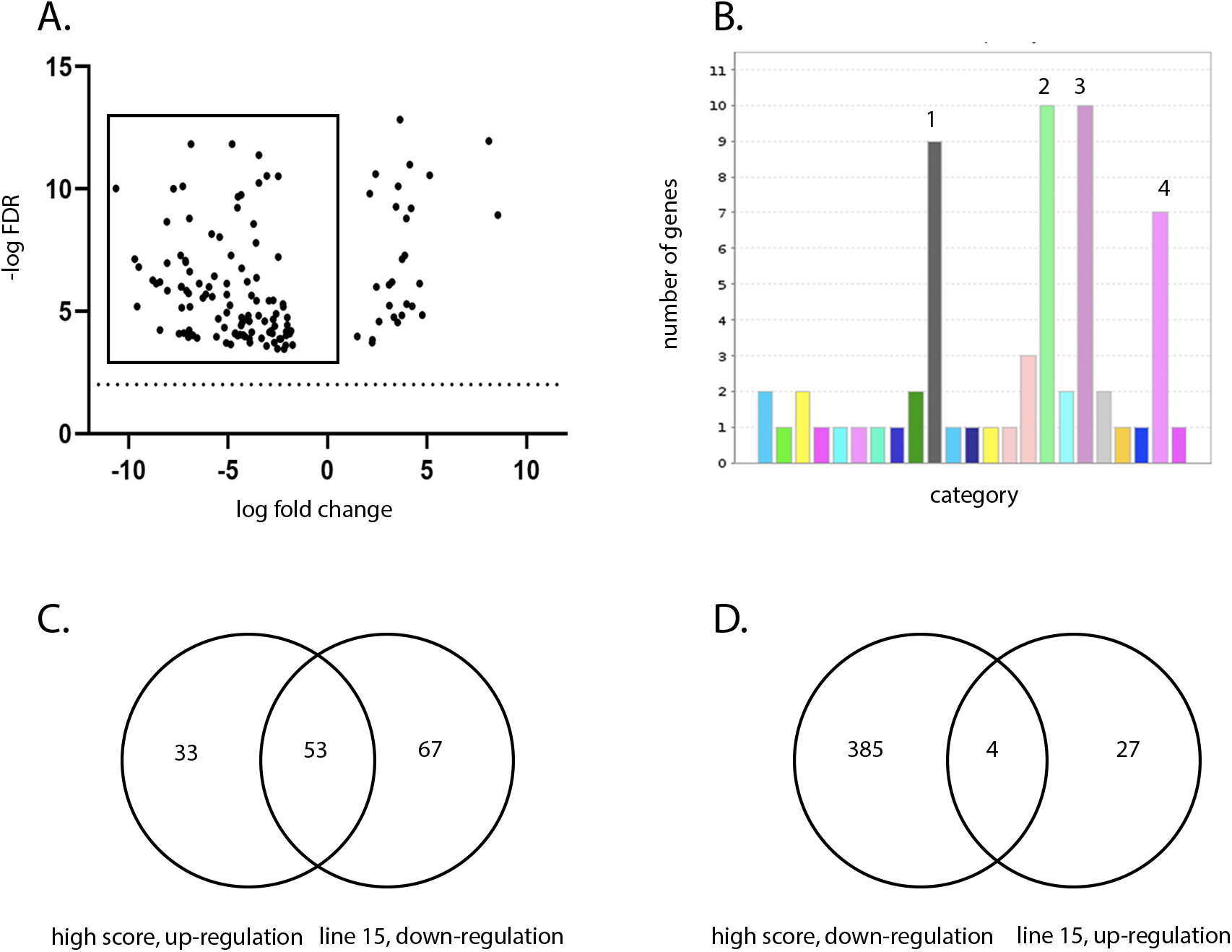
Line W birds showed up-regulation of pathways involved in inflammation mediated by chemokine and cytokine signaling, cytoskeletal regulation by Rho GTPases, nicotinic acetylcholine receptor signaling, and Wnt signaling in the BF compared to line 15 birds. Transcriptional profiles of infected birds at 3 dpi from line W and line 15 were compared, with line W as a baseline. RNA was extracted from the BF of the birds and subjected to RNA-seq. The expression of the target genes was reported as fold-change (FC), with values ≥ 2 considered as up-regulated, and values ≤ -2 considered as down-regulated. The FDR was calculated and a volcano plot of –log FDR over log_2_ FC plotted; FDR values of ≤0.05 were considered as significant. Each dot represents one gene, and genes that were down-regulated in line 15 birds compared to line W (i.e. up-regulated in line W compared to line 15) are boxed (A). Pathway analysis using PANTHER revealed that 4 pathways were significantly up-regulated: 1 (grey)-inflammation mediated by chemokine and cytokine signaling, 2 (green)-cytoskeletal regulation by Rho GTPases, 3 (purple)-nicotinic acetylcholine receptor signaling, and 4 (pink)-Wnt signaling (B). A Venn diagram was plotted of the genes that were up-regulated in birds that had a high clinical score compared to low clinical score and genes that were down-regulated in line 15 birds compared to line W (i.e. genes that were up-regulated in line W birds compared to line 15) (C). A Venn diagram was plotted of the genes that were down-regulated in birds that had a high clinical score compared to low clinical score and genes that were up-regulated in line 15 birds compared to line W (i.e. genes that were down-regulated in line W birds compared to line 15) (D).

### Kinetics of pro-inflammatory gene expression in stimulated primary bursal cells

We determined the kinetics of pro-inflammatory responses in primary cells isolated from the BF of line W and line 15 birds following *ex vivo* infection with IBDV or stimulation with lipopolysaccharide (LPS), a known agonist of pro-inflammatory responses. Primary bursal cells isolated from line 15 birds showed a gradual increase in the expression of pro-inflammatory cytokine genes from 3-18 hours post-IBDV infection, from an average of 1.3 to 2.9 fold IL-1β expression and from 0.6 to 2.2 fold IL-8 expression (Figure 6A and C). The same pattern was observed following LPS stimulation, from an average of 0.7 to 6.3 fold IL-1β expression and from 0.6 to 2.1 fold IL-8 expression 3 -18 hours post-stimulation (Figure 6E and G). In contrast, the kinetics of pro-inflammatory cytokine gene expression differed in cells from line W: Three hours post-LPS stimulation, the average expression of IL-1β was significantly higher in cells isolated from line W birds than line 15 birds (4.0 fold (W) compared to 0.7 fold (15) (p<0.05)), but by 6 hours post stimulation, expression was higher in cells from line 15 birds (Figure 6E). The same trend was observed for IBDV infection, and for IL-6 and IL-8 expression. The expression of inducible nitric oxide synthase (iNOS) also increased following LPS stimulation of cells isolated from line W over the course of the experiment, from an average of 1.0 to 2.2 fold 3-18 hours post-stimulation, but there was no increase in cells from line 15 (Figure 6H). Taken together, these data suggest that primary bursal cells isolated from birds of the more susceptible line W had a more rapid induction of pro-inflammatory responses than cells isolated from the more resistant line 15 following *ex vivo* stimulation.

**Figure 6.**
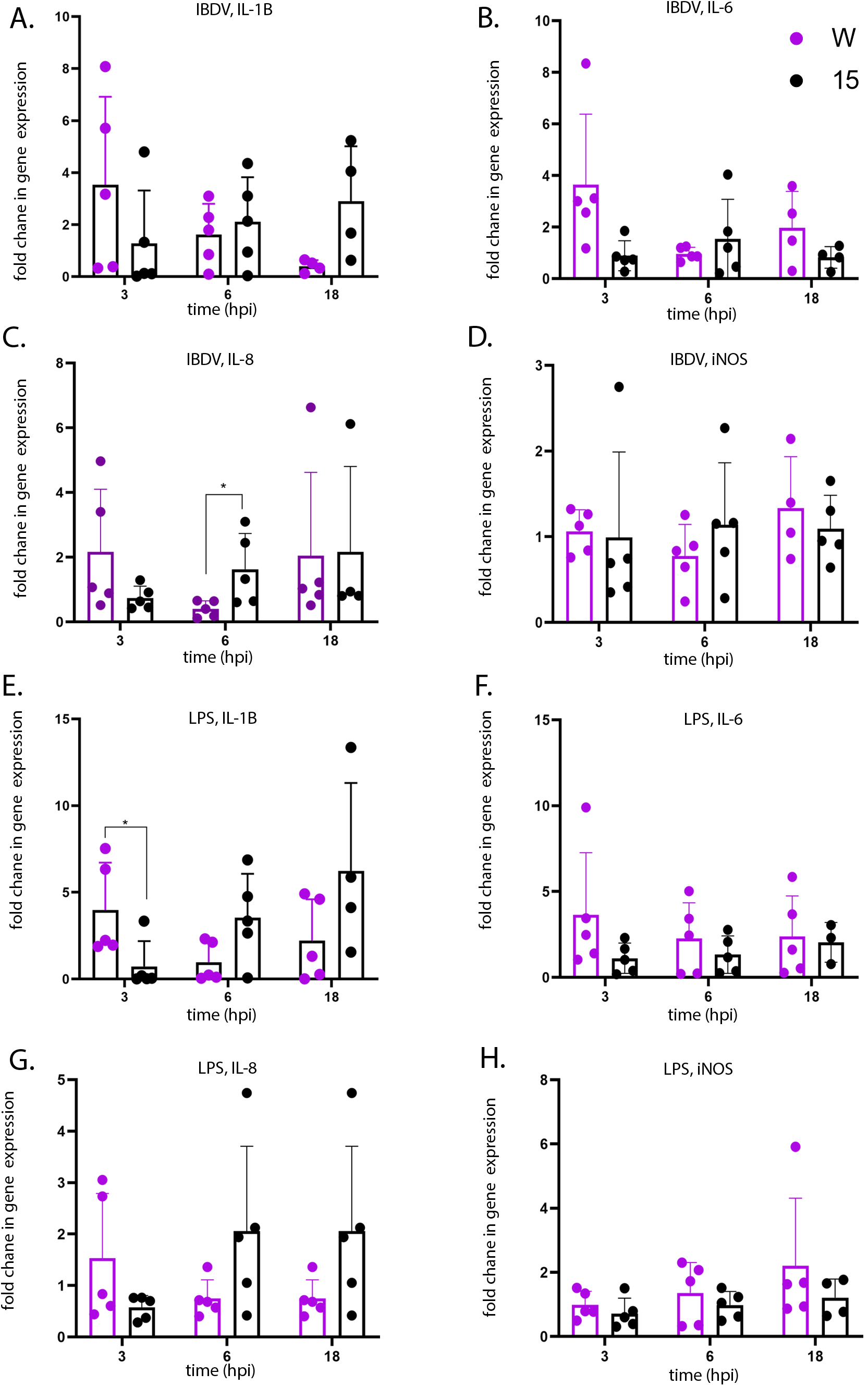
Primary bursal cells isolated from uninfected line W birds and stimulated with LPS *ex vivo* showed a more rapid up-regulation of pro-inflammatory cytokine gene expression compared to cells isolated from uninfected line 15 birds. Primary bursal cells were isolated from the BF of 5 uninfected line W birds and 5 uninfected line 15 birds. The cells were either infected *ex vivo* with IBDV strain UK661 (MOI 3) (A-D) or stimulated *ex vivo* with LPS (100 ng/mL) (E-H). At the indicated time-point post-infection or stimulation, RNA was extracted from the cells, and subjected to RTqPCR. The fold change in the expression of IL-1β (A and E), IL-6 (B and F), IL-8 (C and G) and iNOS (D and H) was quantified, normalized to the housekeeping gene RPLPO and expressed relative to mock-infected or mock-stimulated controls at the same time points in a ΔΔCT analysis. Data passed Shapiro-Wilk normality tests before being analysed by one-way ANOVA and Tukey’ s multiple comparison tests (*p<0.05).

### Quantitation of KUL01+ cells in experimentally inoculated inbred lines

Macrophages are a major source of pro-inflammatory cytokines. We therefore quantified the percentage of KUL01+ macrophage cells and Bu1+ B cells in the bursal cell populations isolated from five uninfected line W and five uninfected line 15 birds by flow cytometry. While there was no significant difference in the percentage of Bu1+ cells (Figure 7A), cells isolated from the BF of uninfected birds from line W contained a significantly higher percentage of KUL01+ cells than cells isolated from line 15 birds (2.8% (W) compared to 1.9% (15) (p<0.01)) (Figure 7B).

**Figure 7.**
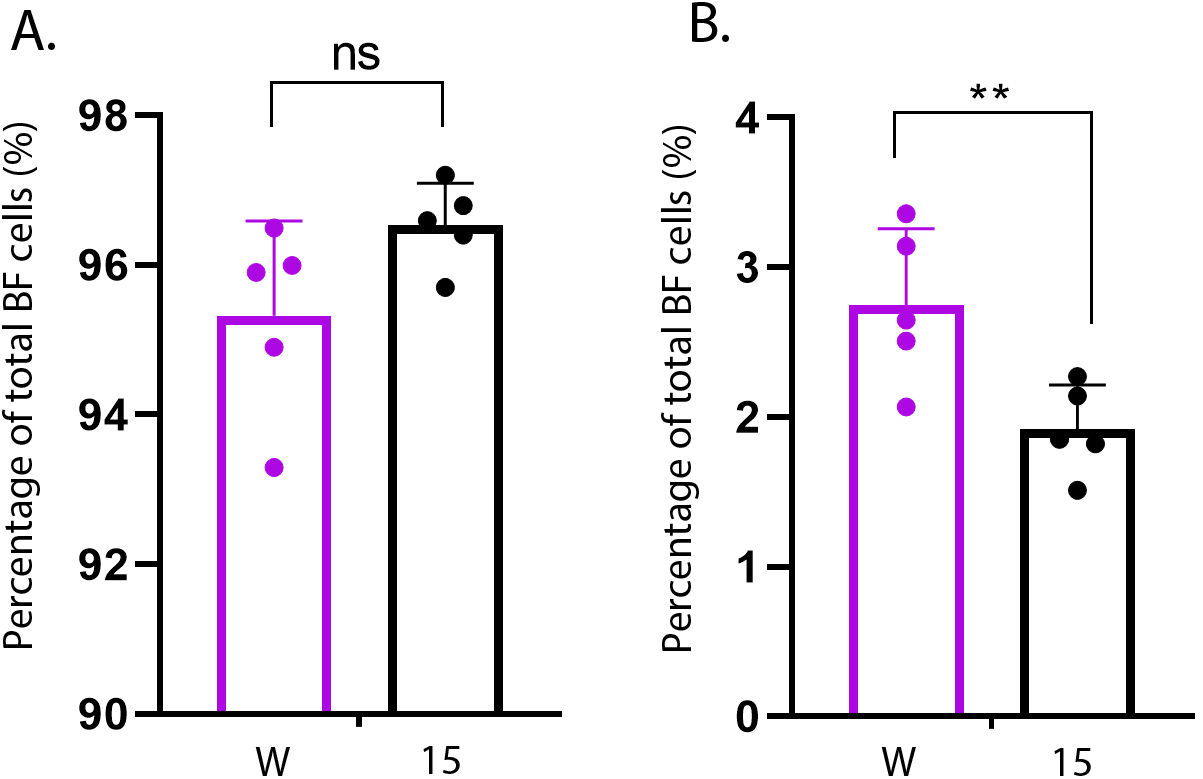
Bursal cells isolated from line W birds contained a greater proportion of KUL01+ cells than cells isolated from line 15 birds. Primary bursal cells were isolated from the BF of uninfected line W and line 15 birds (n=5). Cells were stained with mouse monoclonal antibodies raised against either the B cell marker Bu1, or the macrophage cell marker KUL01. Cells were subjected to flow cytometry and the proportions of Bu1+ cells and KUL01+ cells in the BF cell populations were determined (shown as percentages in A and B, respectively). Data passed Shapiro-Wilk normality tests before being analysed by one-way ANOVA and Tukey’ s multiple comparison tests (**p<0.01).

## Discussion

There is an incentive to understand the molecular basis for differences in disease outcome among inbred lines of White Leghorn chickens infected with IBDV, in order to engineer or breed more disease resistant layer birds that suffer from fewer economic losses. To address this, we selected line W birds as most susceptible, to compare with lines 15, C, and O, previously categorised as “resistant” to IBD (12) with no mortality (7). Birds in all the inbred lines inoculated with vv IBDV strain UK661 became infected and showed clinical signs and bursal lesions. This is consistent with results of Farhanah and Mohd *et al* (10, 11), who showed that all inbred White Leghorn lines succumbed to disease following inoculation with 5.2 log_10_ EID_50_/bird. We reasoned that this high dose may have masked differences between the inbred lines, so we inoculated birds with a lower dose of virus (1.2 log_10_ EID_50_/bird). Even though the birds were inoculated with the same strain and dose of IBDV and were genetically inbred, there was a large variation in the data; for example, clinical scores ranged from 1 to 14 and bursal pathology scores ranged from 0 to 5. We speculate this may have been the result of using a lower dose of inoculum. Despite this variation, we observed that line 15 birds experienced less severe disease compared to other lines, consistent with the data of Farhanah *et al*, who found that lines N and 15 had the lowest bursal lesion scores and lowest virus titre out of six lines tested. Taken together, our data and those of others therefore demonstrate that individual birds of “resistant” inbred lines are not all resistant to IBDV infection.

We compared the transcriptional profile of the BF of birds from line W and line 15 at 3 days post IBDV infection by RNA-seq. As some birds within each experimental group experienced more severe disease and some experienced only mild disease, we also compared the BF transcriptional profile between birds with a high clinical score and birds with a low clinical score, irrespective of the line. RNA-seq analysis revealed that the severity of IBD was associated with up-regulation of pathways involved in inflammation mediated by chemokine and cytokine signaling, cytoskeletal regulation by Rho GTPases, nicotinic acetylcholine receptor signaling, and Wnt signaling in the BF. Interestingly, the same pathways were up-regulated in line W versus line 15 birds and in high clinical score versus low clinical score, and in birds where there was no significant difference in the level of virus replication.

The observation that birds more susceptible to severe disease had an up-regulation in inflammation pathways is consistent with the known literature: Aricibasi *et al* demonstrated that layer-type birds had more severe disease than broilers, with increased expression of Type I IFN, IL-6, IL-1B and IFNγ in the BF (3), and Ruby *et al* found that the White Leghorn line 6 had higher expression of IL-6, IL-8, iNOS, MMP13, IFNα, IFNγ, and cathespin D at 24 hpi than the Brown Leghorn breed by microarray (8). Among genetically-defined, inbred lines within the White Leghorn breed, Mohd *et al* showed by RNA-seq that Line P, which suffered from worse bursal scores and higher viral replication than line N, had increased expression of IL-1β, IL-6, IL-18, IL-17, IL-12B, IFNβ, IFNγ, STAT1 and IL-2 (11). It is known that during IBDV infection there is an influx of KUL01+ macrophages into the BF, and it has been suggested that these infiltrated cells are responsible for the elevated pro-inflammatory cytokine production (4).

Immune cell extravasation into the BF may also be responsible for our observation that cytoskeletal regulation by Rho GTPases and Wnt signaling pathways were up-regulated in birds more susceptible to severe disease as Rho family GTPase signaling is responsible for actin cytoskeleton remodeling that is involved in the migration of immune cells during inflammation, as well as phagocytosis in macrophages (24, 25). Moreover, in the presence of WNT proteins secreted from endothelial cells, T cells produce matrix metalloproteinases that degrade the extracellular matrix, permitting extravasation. Macrophages are also a source of WNT proteins, with classically activated macrophages expressing WNT5A that induces IL-12 production and the consequent production of IFNγ from Th1 cells (26, 27) and alternatively activated macrophages expressing WNT proteins that regulate fibrosis (28). The up-regulation of nicotinic acetylcholine receptor signaling could also be related to inflammation, and recent studies have identified an inflammatory reflex where stimulation of the parasympathetic nervous system by inflammatory mediators leads to increased secretion of acetyl choline (ACh) that binds to nicotinic ACh receptors on macrophages, reducing inflammation (29). Many of the up-regulated genes that were involved in these pathways involved myosin genes. Myosin 1 proteins contain a single heavy chain, are monomeric, and do not form filaments. In B lymphocytes, they regulate cytoskeleton plasticity, cell migration, exocytosis and endocytosis (30). In phagocytic cells, myosin light chain kinase regulates actino-myosin reorganization that underpins differentiation of a monocyte to a macrophage (31). Therefore, the up-regulation of these genes could also be explained by bursal inflammation.

Pro-inflammatory cytokine responses very early following infection of the BF are likely responsible for determining the extent of immune cell extravasation and the degree of inflammation at later time points. Therefore, in order to determine whether there were differences in the kinetics of the early pro-inflammatory cytokine responses between the lines, we isolated primary bursal cells from uninfected line 15 and W birds and either infected them *ex vivo* with IBDV, or stimulated them with LPS. There was a significantly greater expression of IL-1β in cells isolated from line W birds at 3 hours post-stimulation compared to cells isolated from line 15 birds, and the same trend was also true for IBDV infection and for IL-6 and IL-8 expression, suggesting that bursal cells from line W birds responded to IBDV infection and LPS stimulation with a more rapid pro-inflammatory response than cells from line 15 birds. Moreover, this correlated with the percentage of KUL01+ macrophage-like cells in the primary bursal cell population. Taken together, these data are consistent with a model where the BF of more susceptible line W birds contain a higher proportion of macrophages than the more resistant line 15 birds, leading to a more rapid induction of pro-inflammatory responses early following IBDV infection, leading to more bursal immunopathology and more severe clinical disease at later time points.

A direct comparison of line W and 15 birds has previously been conducted following *Salmonella* infection, where line W was found to be more resistant than line 15 to *S. typhimurium* (32). In another study, primary macrophages isolated from the more resistant line W birds and infected *ex vivo* with *S. enterica* and *S. Gallinarum* showed a more rapid pro-inflammatory response to a greater level than cells from the more susceptible line 7 birds (33). This implies that a more rapid and robust pro-inflammatory response may be a feature of cells from line W birds and, while this may be protective against *Salmonella* infections, it may lead to excessive inflammation and pathology in IBDV infection. The reason why cells from line W birds may induce more rapid pro-inflammatory responses to IBDV infection, *Salmonella* infection or LPS stimulation, compared to other lines, remains unknown, and it is possible that single nucleotide polymporphisms (SNPs) in genes of the pro-inflammatory pathway are responsible for differences in the kinetics of the responses. Alternatively, SNPs in the promoter regions or regulatory elements, and possibly also epigenetic changes in genes of the pro-inflammatory pathway may also be responsible for the observed differences, and more work is needed to explore this further.

While our data suggest that inflammation is a major factor in determining the severity of IBD, there may be additional mechanisms that contribute to bursal pathology and disease outcome. For example, it is known that a strong T cell response in the BF correlates with lower disease severity (3). An influx of T cells into the BF could be responsible for the transcriptional profile we observed *in vivo*, and it is possible that impaired or inappropriate T cell responses may contribute to the pathology seen in more susceptible birds compared to more resistant birds. The inbred lines differ in their Major Histocompatility Complex (MHC) haplotypes, which are responsible for presenting peptide antigens to T cells. Polymorphisms within the MHC gene between the different lines can influence the disease outcome of Marek’ s disease and avian leukosis (34, 35), and it is possible that the same might be true for IBDV. Mohd *et al* observed that haplotype B21 (line N) was more resistant than haplotype B19 (line P) (11). We extend this observation by finding that haplotype B15 (line 15) was also more resistant than haplotype B14 (line W) but a characterisation of the T cell responses was beyond the scope of this study, as we only considered time points up to 3 dpi, and T cell influx into the BF does not peak until later (36).

Interestingly, we noted that interferon stimulated genes (ISGs) were not up-regulated in high versus low clinical score or in line 15 versus W birds. However, it should be noted that RNA-seq was only performed using samples from infected birds, and we did not compare the gene expression between mock and infected birds. It is likely that ISGs were up-regulated in infected birds relative to mock as has previously been shown (21). We also observed genes that were up-regulated in line 15 birds compared to line W were involved in a variety of pathways, but no pathway was more represented than another, suggesting that a more balanced transcriptional response existed in the BF of birds that are more resistant to severe disease.

Several studies have now analysed transcriptomic responses between birds of different inbred lines (8-11). In the future, it will be important to consider an analysis of proteomics, given that mRNA expression does not always correlate with protein expression. Moreover, Quantitative Trait Locus (QTL) mapping and Genome Wide Association Studies (GWAS) are also needed to identify genes associated with disease resistance, as have already been done for an aquatic birnavirus, infectious pancreatic necrosis virus (IPNV), in Atlantic salmon and rainbow trout (37, 38).

In summary, among inbred lines of the White Leghorn breed, we found that an up-regulation of inflammatory pathways correlated with enhanced bursal pathology and symptoms *in vivo*. Moreover, we found that primary bursal cells cultured from a susceptible line showed more rapid induction of proinflammatory reponses than primary cells cultured from a more resistant line when stimulated *ex vivo*, which correlated with the number of KUL01+ macrophages in the cell population. We propose a model where more rapid induction of pro-inflammatory responses early following IBDV infection in more susceptible lines exacerbates the inflammation at later time points and leads to more bursal immunopathology and more severe clinical disease relative to more resistant lines. While our data suggest that differences in the inflammatory response between different chicken lines could be due to differences in the number of KUL01+ macrophages in the BF, there may also be differences in the pro-inflammatory pathways between the lines that still need to be determined. Once these differences have been characterised, it may be possible for them to be exploited to engineer a White Leghorn line that suffers from lower production losses due to IBDV. However, if such birds were to be produced, care should be taken to ensure they are not more susceptible to other infections.

## Acknowledgments

AA was supported through the National Centre for the Replacement, Refinement & Reduction of Animals in Research (NC3Rs), grant NC/R001138/1. AB was supported through The Houghton Trust (Small Project Research Grant awarded in 2015), and the BBSRC (grant BBS/E/I/00001845). EG was supported through grant 104771/Z/14/. The Pirbright Institute facilities used in this project were strategically funded through the BBSRC (grants BBS/E/I/00007031 and BBS/E/I/00007038). The funders had no role in study design, data collection and interpretation, or the decision to submit the work for publication.

## Author contributions

AA performed experiments, analyzed data and wrote the first draft of the manuscript; SN, EC and KD performed experiments and edited drafts of the manuscript; VR, EG and MS analyzed data and edited drafts of the manuscript; AB conceptualized the study, obtained funding, helped to conduct experiments and analyze data, and edited drafts of the manuscript.

## Ethics statement

All animal procedures were conducted following the approval of the Animal Welfare and Ethical Review Board (AWERB) at The Pirbright Institute, under Home Office Establishment, Personal and Project licenses, and conformed to the United Kingdom Animal (Scientific Procedures) Act (ASPA) 1986.

## Supplementary Material

**Table S1**. Genes that were up-regulated in birds with a high clinical score compared to birds with a low clinical score, irrespective of the line.

**Table S2**. Genes that were down-regulated in birds with a high clinical score compared to birds with a low clinical score, irrespective of the line.

**Table S3**. Genes that were up-regulated in line 15 birds compared to line W.

**Table S4**. Genes that were down-regulated in line 15 birds compared to line W.

**Table S5**. Primers used in the study

