## Supplementary Table 5 for "Transcriptomic analysis reveals that severity of infectious bursal disease in White Leghorn inbred chicken lines is associated with greater bursal inflammation *in vivo* and more rapid induction of pro-inflammatory responses in primary bursal cells stimulated *ex vivo*"

| **Name** | **Forward Sequence**  **(5’-3’)** | **Reverse Sequence**  **(5’-3’)** |
| --- | --- | --- |
| IBDV | GAGGTGGCCGACCTCAACT | GCCCGGATTATGTCTTTGAAG |
| IL-1β | GCTCTACATGTCGTGTGATGAG | TGTCGATGTCCCGCATGA |
| IL-6 | AACATGCGTCAGCTCCTGAAT | TCTGCTAGGACTTCTCCATTGAA |
| IL-8 | GCCCTCCTCCTGGTTTCAG | TGGCACCGCAGCTCATT |
| iNOS | CCTGGAGGTCCTGGAAGAGT | CCTGGGTTTCAGAAGTGGC |

**Table S5.** Primers used in this study
